# Coherent natural scene structure facilitates the extraction of task-relevant object information in visual cortex

**DOI:** 10.1101/2020.12.01.406959

**Authors:** Daniel Kaiser, Greta Häberle, Radoslaw M. Cichy

**Author notes:** <u>Correspondence:</u> Dr. Daniel Kaiser, Department of Psychology, University of York, Heslington, York, YO10 5DD, UK.

## Abstract

Looking for objects within complex natural environments is a task everybody performs multiple times each day. In this study, we explore how the brain uses the typical composition of real-world environments to efficiently solve this task. We recorded fMRI activity while participants performed two different categorization tasks on natural scenes. In the object task, they indicated whether the scene contained a person or a car, while in the scene task, they indicated whether the scene depicted an urban or a rural environment. Critically, each scene was presented in an “intact” way, preserving its coherent structure, or in a “jumbled” way, with information swapped across quadrants. In both tasks, participants’ categorization was more accurate and faster for intact scenes. These behavioral benefits were accompanied by stronger responses to intact than to jumbled scenes across high-level visual cortex. To track the amount of object information in visual cortex, we correlated multi-voxel response patterns during the two categorization tasks with response patterns evoked by people and cars in isolation. We found that object information in object- and body-selective cortex was enhanced when the object was embedded in an intact, rather than a jumbled scene. However, this enhancement was only found in the object task: When participants instead categorized the scenes, object information did not differ between intact and jumbled scenes. Together, these results indicate that coherent scene structure facilitates the extraction of object information in a task-dependent way, suggesting that interactions between the object and scene processing pathways adaptively support behavioral goals.

## 1 Introduction

Despite the complexity of our everyday environments, perceiving objects embedded in natural scenes is remarkably efficient. This efficiency is illustrated by studies that require participants to categorize objects under conditions of limited visual exposure: For instance, participants can tell whether a scene contains an animal or not from just a single glance (Thorpe et al., 1996; Potter, 1975, 2012), and even when only limited attentional resources are available (Li et al., 2002).

The ability to effortlessly make such categorization responses is underpinned by the efficient extraction of object information in visual cortex. Neuroimaging research has shown that the category of task-relevant objects can be reliably decoded from fMRI activity patterns in visual cortex, even when the objects are embedded in complex natural scenes (Peelen et al., 2009; Peelen & Kastner, 2011; Seidl et al., 2012) or movies (Cukur et al., 2013; Nastase et al., 2017; Shahdloo et al., 2020). M/EEG studies demonstrate that object category is represented well within the first 200ms of vision, even when the object is shown under such naturalistic conditions (Cauchoix et al., 2014; Kaiser et al., 2016; VanRullen & Thorpe, 2001; Thorpe et al., 1996). Together, these results highlight that the cortical processing of objects appearing within rich real-world environments is surprisingly efficient.

This processing efficiency becomes less surprising if scene context is not just considered as a nuisance that puts additional strain on our visual resources. Indeed, contextual information can facilitate object processing (Bar, 2004): For instance, scene context allows for efficient allocation of attention (Torralba et al., 2006; Wolfe et al., 2011; Võ et al., 2019), or for disambiguating object information under uncertainty (Brandmann & Peelen, 2017; Oliva & Torralba, 2007). Such findings demonstrate that object and scene processing mechanisms interact with each other to enable the efficient processing of object information.

Here, we investigated how the coherent spatial structure of the scene context aids the extraction of object information from the scene. To this end, we used a jumbling paradigm, in which we disrupted the scenes’ coherent structure by dividing them into multiple rectangular pieces and shuffling those pieces. Classical studies suggest that jumbling drastically impairs participants’ ability to categorize both the scene itself (Biederman et al., 1974), and the object embedded within the scene (Biederman et al., 1972, 1973). Such impairments can be linked to changes in cortical scene processing: We have recently shown that scene-selective brain responses are less pronounced and contain less scene category information when the scene is jumbled (Kaiser et al., 2020a, 2020b). However, it is unclear how these changes in scene-selective activations modulate the representation of objects within the scene.

In the current study, we thus set out to characterize how the presence of an intact – versus a jumbled – scene context modulates object representations in visual cortex. First, we asked whether cortical object processing is indeed facilitated by the presence of a coherent scene context. Second, we asked whether such facilitation effects depend on the objects being relevant or irrelevant for current behavioral goals.

To answer these questions, we recorded fMRI activity while participants categorized objects contained in intact or jumbled scenes. We found that fMRI responses across high-level visual cortex were generally higher for intact scenes than for jumbled scenes, revealing widespread sensitivity to scene structure. When analyzing object category information in multi-voxel response patterns, we found that coherent scene structure enhanced object information in object-selective visual cortex. However, this enhancement was task-specific: When participants categorized the scenes instead of the objects, we found no such enhancement of object information. These results suggest that the visual brain uses coherent real-world structure to more efficiently extract task-relevant object information from complex scenes.

## 2 Materials and Methods

### 2.1 Participants

Twenty-five healthy adults (mean age 26.4 years, SD=5.3; 15 female, 10 male) participated. All participants had normal or corrected-to-normal vision. They all provided informed written consent and received either monetary reimbursement or course credits. Procedures were approved by the ethical committee of the Department of Psychology at Freie Universität Berlin and were in accordance with the Declaration of Helsinki.

### 2.2 Stimuli

The stimulus set consisted of colored natural scene photographs (640×480 pixels resolution). Scenes were selected to cover three independent manipulations. First, each scene contained one of two object categories: half of the scenes contained a person (or multiple people), whereas the other half contained a car (or multiple cars). Second, the person or car appeared equally often in each of the quadrants of the scene. Third, each scene belonged to one of two scene categories: half of the scenes depicted urban environments, the other half depicted rural environments. For each possible combination of these factors (e.g., a person appearing in the bottom left quadrant of a rural scene), 10 unique scene exemplars were available, yielding 160 scenes in total (2 object categories × 4 object locations × 2 scene categories × 10 exemplars). During the experiment, the scenes could be presented in their original orientation or mirrored along their vertical axis (as in Kaiser et al., 2016), yielding a total of 320 different scene stimuli. Example scenes are shown in Figure 1a.

**Figure 1.**
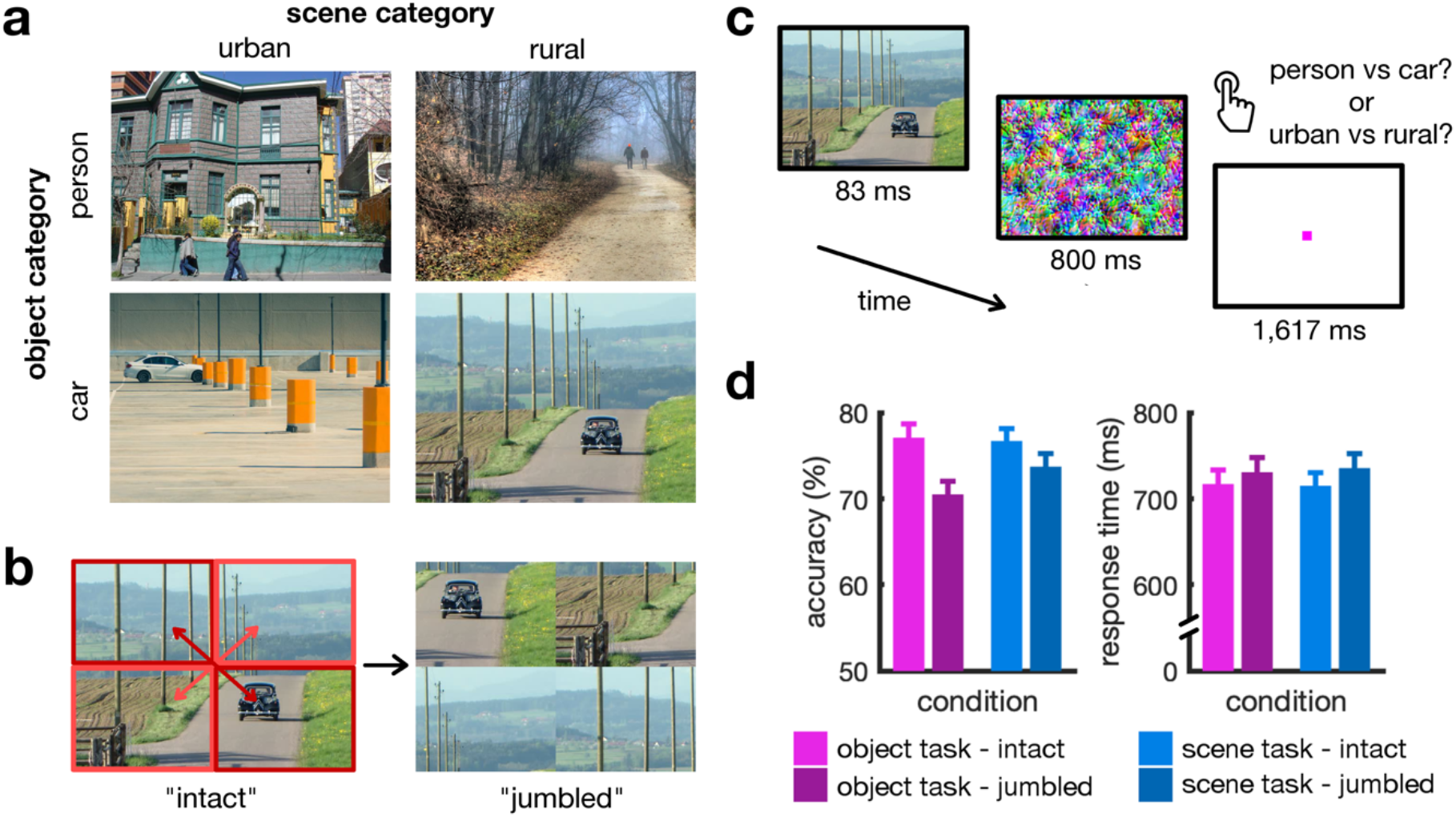
Stimuli, paradigm, and behavioral results. **a)** Stimuli consisted of natural scene images from two categories: urban or rural environments. Each of the scenes contained one of two object categories: people or cars. **b)** During the experiment, these scenes were shown in an unaltered way (“intact” condition) or with their quadrants intermixed (“jumbled” condition). The jumbled scenes were created by shuffling the quadrants in a crisscrossed way, as illustrated. **c)** Participants viewed each scene briefly, followed by a visual mask. In separate runs, they performed two different tasks: They were either asked to indicate whether the scene contained a person or a car (“object task”) or whether the scene depicted an urban or a rural environment (“scene task”). **d)** In both tasks, scene structure impacted behavioral performance: Participants were significantly more accurate and faster for the intact scenes than for the jumbled scenes. Error bars represent standard errors of the mean.

To manipulate scene structure, we either presented the scenes in a coherent, “intact” condition or in an incoherent, “jumbled” condition. Jumbled scenes were generated by shuffling the four quadrants of the image in a crisscrossed way (i.e., top-left was swapped with bottom-right, and top-right was swapped with bottom-left; Figure 1b). This manipulation solely affected the scene’s structure, but not the people or cars contained in the scene: First, as the objects never straddled the boundary between quadrants, the objects themselves always remained unaltered. Second, as the objects appeared equally often in each quadrant before jumbling the scenes, they also appeared equally often in each quadrant after jumbling them.

In total, 640 scene images were used, which covered 320 intact scenes and 320 jumbled scenes. Additionally, 200 colored texture masks (Kaiser et al., 2016) were used to visually mask the scenes during the experiment (see below).

### 2.3 Experimental Paradigm

Each participant completed four experimental runs of 17 minutes each. Each run contained 320 experimental trials, corresponding to 320 unique scene stimuli. Both intact and jumbled scenes were included in each run. For half of the participants, the even runs only contained the original scenes, while the odd runs only contained the horizontally mirrored scenes; for the other half of the participants, the odd runs only contained the original scenes, while the even runs only contained the horizontally mirrored scenes. Each of the scenes was presented once during the run. Trial order was fully randomized for each participant and run.

On each trial, the scene was presented for 83ms, immediately followed by a visual mask (chosen randomly from the 200 available masks) for 800ms. Masks were shown to establish a sensitive performance range for reasonably long presentation times, as they disrupt ongoing visual processing after the offset of the stimulus. All images were shown within a black rectangle (10deg X 7.5deg visual angle). After an inter-trial interval of 1,617ms, during which a pink fixation dot was shown, the next trial started. An example trial is illustrated in Figure 1c. In addition to the experimental trials, each run contained 80 fixation-only trials, during which only the fixation dot was displayed. Runs started and ended with a brief fixation period.

In two of the four runs, participants were asked to categorize the object contained in each scene as either a person or a car (“object task”). In the other two runs, participants were asked to categorize the scene as either a rural or an urban environment (“scene task”). Participants were instructed to respond as accurately and quickly as possible, with an emphasis on accuracy. Button-press responses were recorded during the whole inter-trial interval (i.e., until 2,500s after stimulus onset). The four runs were alternating between the object and scene tasks. The task in the first run was counter-balanced between participants. Notably, physical stimulation was completely identical across the object and scene tasks.

All stimuli were back-projected onto a translucent screen mounted to the head end of the scanner bore. Participants viewed the stimulation through a mirror attached to the head coil. Stimulus presentation was controlled using the Psychtoolbox (Brainard, 1997).

### 2.4 Benchmark Localizer Paradigm

In addition to the experimental runs, each participant completed a benchmark localizer run, which was designed to obtain “benchmark” patterns in response to people and cars in isolation (Peelen et al., 2009; Peelen & Kastner, 2011). During this run, participants viewed images of bodies, cars, and scrambled images of bodies and cars. For each of the three categories, 40 images were used. All images were different than the ones used in the main experiment. These images were presented in a block design. Each block lasted 20 seconds and contained 20 images of one of the three categories, or only a fixation cross. Images were presented for 500ms (5deg × 5deg visual angle), separated by a 500ms inter-stimulus interval. The benchmark localizer run consisted of a total of 24 blocks (6 blocks for each of the three stimulus categories, and 6 fixation-only blocks). Four consecutive blocks always contained the four different conditions in random order. Participants were instructed to respond to one-back image repetitions (i.e., two identical images back-to-back), which happened once during each non-fixation block. The benchmark localizer run lasted 8:30 minutes and was completed halfway through the experiment, after two of the four experimental runs.

### 2.5 fMRI recording and preprocessing

MRI data was acquired using a 3T Siemens Magnetom Tim Trio Scanner equipped with a 12-channel phased-array head coil. T2*-weighted gradient-echo echo-planar images were collected as functional volumes, with the following parameters: TR=2s, TE=30ms, 70° flip angle, 3mm3 voxel size, 37 slices, 20% slice gap, 192mm FOV, 64×64 matrix size, interleaved acquisition, A/P phase encoding, acquisition time 17min (main experiment) / 8:20min (benchmark localizer), whole-brain coverage, ACPC orientation. Additionally, a T1-weighted 3D MPRAGE image was obtained as an anatomical reference, with the following parameters: TR=1.9s, TE=2.52ms, 9° flip angle, 1mm3 voxel size, 176 slices, 50% slice gap, 256mm FOV, ascending acquisition, A/P phase encoding, acquisition time 4:26min, whole-brain coverage. All acquisitions contained four initial dummy volumes that were discarded later.

Preprocessing and hemodynamic response modelling was performed using SPM12 (www.fil.ion.ucl.ac.uk/spm/). Functional volumes were realigned and coregistered to the anatomical image. Further, transformation parameters to MNI-305 standard space were obtained using the “segmentation” routine in SPM12.

Functional data from each experimental run were modelled in a general linear model (GLM) with 16 experimental predictors (2 object categories × 4 object locations × 2 scene categories). Additionally, we included the six movement regressors obtained during realignment. Data from the benchmark localizer run were modelled in a GLM with three experimental predictors (person, car, scrambled) and six movement regressors.

### 2.6 Region of interest definition

We restricted fMRI analyses to five regions of interest (ROIs): early visual cortex (EVC), object-selective lateral occipital cortex (LO), body-selective extrastriate body area (EBA), scene-selective occipital place area (OPA), and scene-selective parahippocampal place area (PPA). ROIs masks were defined using group-level activation masks from functional brain atlases: For EVC, we selected all voxels that were most probably assigned to primary visual cortex (V1v, V1d) in the Wang et al. (2015) atlas, and for LO, EBA, OPA, and PPA we selected region masks from the Julian et al. (2012) atlas. ROIs were defined separately for each hemisphere. All ROI masks were inverse-normalized into individual-participant space using the parameters obtained during T1 segmentation. Average voxel counts in individual-participant space amounted to 248/271 (EVC; SD=42/41, left/right), 929/947 (LO; SD=103/102), 402/443 (EBA; SD=45/52), 26/47 (OPA; SD=5/8), and 140/105 (PPA; SD=14/10). Notably, the LO and EBA ROIs overlapped to some extent (300/406 voxels overlap, left/right); the inclusion of the EBA allowed us to see whether the results hold in a smaller cortical region with a narrower category preference for bodies. As we did not have any hypothesis related to hemispheric differences, all results for the left- and right-hemispheric ROIs were averaged before statistical analysis. Separate results for the right- and left-hemispheric ROIs are reported in the Supplementary Information.

### 2.7 Univariate analysis

Response magnitudes during the experimental runs were analyzed separately for each ROI. We first averaged beta values across the two object-task and scene-task runs, respectively. We then averaged beta values across object categories, object locations, and scene categories. This way, we obtained response magnitudes for four conditions: (1) responses to intact scenes in the object task, (2) responses to jumbled scenes in the object task, (3) responses to intact scenes in the scene task, and (4) responses to jumbled scenes in the scene task. These four conditions allowed us to separately estimate the effects of task (object task versus scene task) and scene structure (intact versus jumbled) on neural responses across the five ROIs. For a univariate analysis of category-specific responses across the two tasks, see the Supplementary Information.

### 2.8 Multivariate pattern analysis

Multivariate pattern analysis (MVPA) was carried out in CoSMoMVPA (Oosterhof et al., 2016). Our MVPA approach closely followed similar fMRI studies that investigated the representation of objects in natural scenes (Peelen et al., 2009; Peelen & Kastner, 2011). We first computed a one-sample t-contrasts for every condition against baseline (i.e., against the fixation trials). In the benchmark localizer run, there were 2 such t-contrasts (one for people versus baseline, and one for cars versus baseline). In the object task and scene task runs, there were 16 t-contrasts each (one contrast for each experimental condition against baseline, reflecting 2 object categories × 4 object locations × 2 scene categories). For each of the three tasks (benchmark localizer, object task, and scene task), the resulting t-values were normalized for each voxel by subtracting the average t-value across conditions. For each ROI, multi-voxel response patterns were constructed by concatenating the t-values across all voxels belonging to the ROI.

To obtain an index of object discriminability (i.e., how discriminable people and cars in scenes are based on multi-voxel response patterns), we performed a correlation-based MVPA. The goal of this analysis was to quantify how “person-like” or “car-like” the cortical representation of each of the scenes was, thereby isolating the amount of object category information in visual cortex (note that each of the scenes either contained a person or a car). To this end, we correlated multi-voxel response patterns evoked by people and cars in isolation (from the benchmark localizer) with response patterns evoked by people and cars contained in a scene (from one of the experimental tasks). These correlations were Fisher-transformed. To quantify object discriminability, we then subtracted the correlations between different categories (e.g., person in isolation and car within a scene) from correlations between the same categories (e.g., person in isolation and person within a scene). This yielded an index of category-discriminability, with values greater than zero indicating that the two categories are represented differently (Haxby et al., 2001). Results for different analysis routines (using Spearman correlations and no mean-removal across conditions) can be found in the Supplementary Information.

Before performing this analysis, response patterns in the main experiment were averaged across object locations and scene categories. This way, we obtained an index of object category-discriminability for four separate conditions: (1) category-discriminability for intact scenes in the object task, (2) category-discriminability for jumbled scenes in the object task, (3) category-discriminability for intact scenes in the scene task, and (4) category-discriminability for jumbled scenes in the scene task. The resulting four conditions allowed us to estimate the effects of scene structure on the quality of object representations in visual cortex, both when the objects were task-relevant and task-irrelevant.

### 2.9 Statistical testing

To compare behavioral performance, univariate responses, and multi-voxel pattern information across conditions, we used repeated-measures ANOVAs and paired-sample t-tests. We report partial eta-squared (η_p_^2^, for F-tests) and Cohen’s d (for t-tests) as measures of effect size. Descriptive statistics (means and standard errors) are reported in the Supplementary Information.

### 2.10 Data availability

Data are publicly available on OSF (doi.org/10.17605/osf.io/gs2t5). Other materials are available from the corresponding author upon request.

## 3 Results

### 3.1 Coherent scene structure facilitates the perception of objects within scenes

We first analyzed participants’ behavioral performance in the object and scene tasks, separately for the intact and jumbled scenes (Figure 1d). In the object task, participants’ categorization (person versus car) of objects within the intact scenes was more accurate, t(24)=8.28, p<.001, d=1.61, and faster, t(24)=3.26, p=.0033, d=0.65, compared to the jumbled scenes. In the scene task, participants’ categorization (rural versus urban) of the intact scenes was more accurate, t(24)=4.77, p<.001, d=0.95, and faster, t(24)=3.26, p=.0033, d=0.65, compared to the jumbled scenes. These results are in line with classical findings on object and scene categorization in jumbling paradigms (Biederman, 1972; Biederman et al., 1973, 1974), showcasing that scene jumbling has a profound impact on perception.

Further, when directly comparing the two tasks, we did not find differences in accuracy, F(1,24)=3.13, p=.090, or response times, F(1,24)=0.04, p=.84. Any differences in neural responses are therefore unlikely to reflect differences in task difficulty, and therefore attentional engagement, between the two tasks.

Together, these results demonstrate that jumbling similarly impairs the perception of the scene and the objects contained in it, demonstrating a cross-facilitation between scene and object vision that can be observed on the behavioral level.

### 3.2 Scene structure impacts univariate responses across object- and scene-selective cortex

To quantify the effects of scene jumbling on the neural level, we first ran univariate analyses. In these analyses, we compared fMRI response magnitudes across the intact and jumbled scenes and across the two tasks (Figure 2). To do so, we performed a 2×2 repeated measures ANOVA with the factors scene structure (intact versus jumbled) and task (object task versus scene task). The analysis was performed separately and in turn for each of the five ROIs: EVC, LO, EBA, OPA, and PPA. Detailed results for these analyses can be found in Table 1.

**Table 1:** Univariate responses, analyzed in a 2×2 repeated measures ANOVA with the factors scene structure (intact versus jumbled) and task (object task versus scene task). Significant effects are highlighted in bold.

| ROI | Main Effect<br>Scene Structure | Main Effect<br>Task | Interaction Effect<br>Structure × Task |
| --- | --- | --- | --- |
| **EVC** | $F(1,24) < 0.01$, $p = .98$, $\eta_p^2 < 0.01$ | $F(1,24) = 1.25$, $p = .28$, $\eta_p^2 = 0.05$ | $F(1,24) = 0.09$, $p = .76$, $\eta_p^2 < 0.01$ |
| **LO** | **$F(1,24) = 9.74$, $p = .005$, $\eta_p^2 = 0.29$** | $F(1,24) = 0.04$, $p = .85$, $\eta_p^2 < 0.01$ | $F(1,24) = 0.97$, $p = .33$, $\eta_p^2 = 0.04$ |
| **EBA** | **$F(1,24) = 7.95$, $p = .009$, $\eta_p^2 = 0.25$** | $F(1,24) = 0.21$, $p = .65$, $\eta_p^2 < 0.01$ | $F(1,24) = 2.46$, $p = .13$, $\eta_p^2 = 0.09$ |
| **OPA** | **$F(1,24) = 27.18$, $p < .001$, $\eta_p^2 = 0.53$** | $F(1,24) = 0.09$, $p = .77$, $\eta_p^2 < 0.01$ | $F(1,24) = 0.97$, $p = .34$, $\eta_p^2 = 0.04$ |
| **PPA** | **$F(1,24) = 48.02$, $p < .001$, $\eta_p^2 = 0.67$** | **$F(1,24) = 6.51$, $p = .017$, $\eta_p^2 = 0.21$** | $F(1,24) = 0.51$, $p = .48$, $\eta_p^2 = 0.02$ |

**Figure 2.**
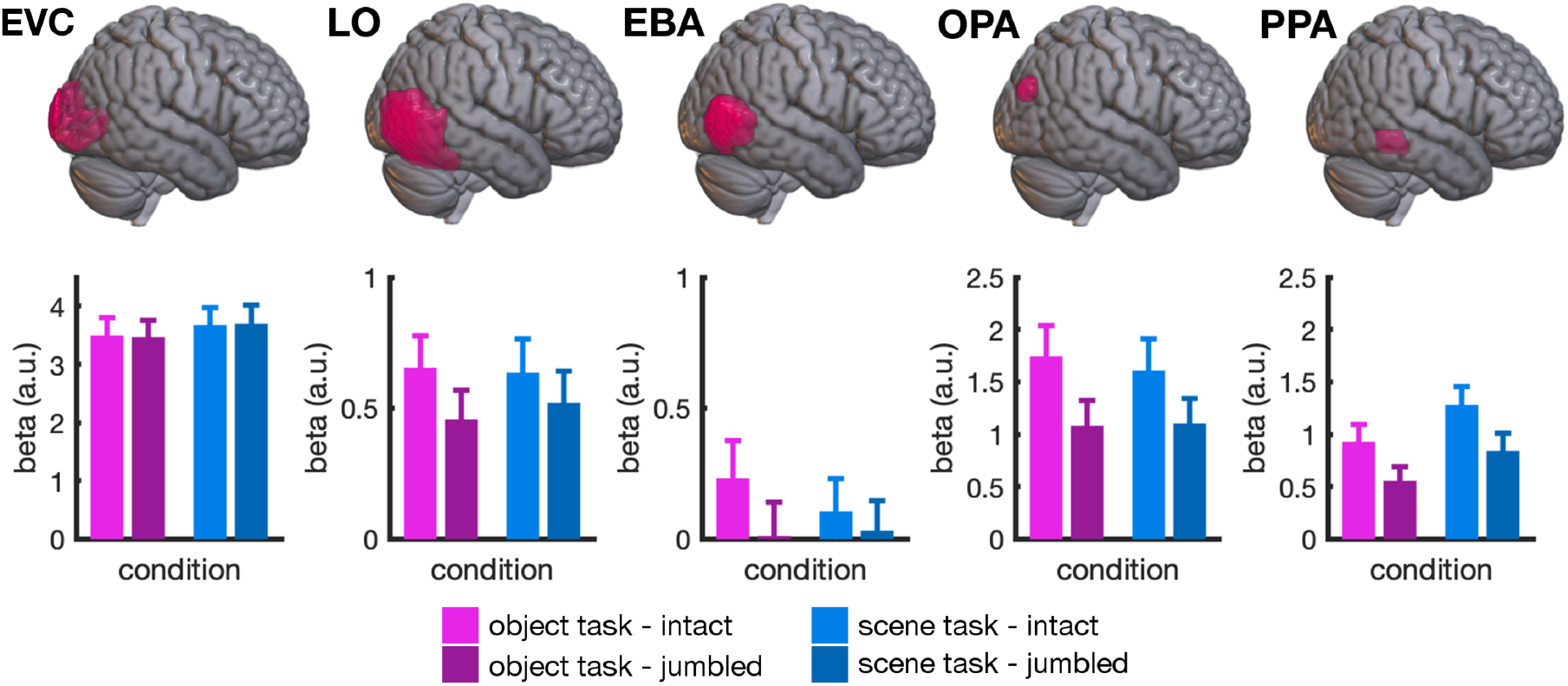
Univariate results. In all extrastriate regions, but not in EVC, we found a significant main effect of scene structure: Intact scenes led to significantly stronger responses than jumbled scenes. This effect was comparable across the two tasks and most pronounced in scene-selective ROIs. PPA was the only region that additionally showed a modulation by task, with significantly stronger responses when participants were categorizing the scenes, compared to when they were categorizing the objects within them. For illustration purposes, ROI masks are shown on the right hemisphere of a standard-space template using MRIcroGL (Li et al., 2016); the displayed results are averaged across ROIs in both hemispheres. Error bars represent standard errors of the mean.

In EVC, responses were comparable across all conditions, all F<1.25, p>.27, η_p_^2^<0.06, suggesting that EVC is not sensitive to typical scene composition.

In all extrastriate ROIs, we found a main effect of scene structure, which indicated stronger responses to intact than to jumbled scenes, all F(1,24)>7.95, p<.010, η_p_^2^>0.24. Comparing this effect across regions, we found that it was more pronounced in the scene-selective regions, OFA versus LO/EBA, both F(1,24)>31.17, p<.001, η_p_^2^>0.56, and PPA versus LO/EBA, both F(1,24)>35.54, p<.001, η_p_^2^>0.59. This finding confirms our previous fMRI results, which revealed particularly strong effects of scene jumbling in scene-selective areas of visual cortex (Kaiser et al., 2020a).

In all ROIs, scene structure affected univariate responses similarly across the two tasks, as indexed by no significant interaction effects, all F<2.46, p>.12, η_p_^2^<0.10. This pattern of results mirrors the pattern observed in behavior, where scene jumbling produced comparable effects in the object and scene tasks.

PPA was the only region that additionally showed an effect of task, F(1,24)=6.51, p=.017, η_p_^2^=0.21, with stronger responses in the scene task compared to the object task. This suggests an increased importance of computations in higher-level scene-selective cortex when scene attributes were behaviorally relevant.

Having established that scene structure enhanced cortical responses across object- and scene-selective cortex, and similarly for both tasks, we next asked how scene structure contributes to the extraction of object information – both when the objects are behaviorally relevant and when they are not.

### 3.2 Coherent scene structure enhances task-relevant object information in multi-voxel response patterns

To understand how the coherent spatial structure of the scene impacts cortical object processing, we performed a correlation-based multivariate pattern analysis (MVPA). In this analysis, we correlated the multi-voxel response patterns evoked by objects embedded in scenes (from the object and scene tasks) with the patterns evoked by the objects in isolation (from the benchmark localizer) (Figure 3a). This approach allowed us to quantify how “person-like” or “car-like” the cortical representation of each of the scene conditions was, thereby isolating the amount of object information present in visual cortex (note that each of the scenes either contained a person or a car). When object information is operationalized in this way, it can be separated from differences in the scene context (as in the benchmark localizer no scene context is presented) and task-related differences (as in the benchmark localizer participants perform a different task).

**Figure 3.**
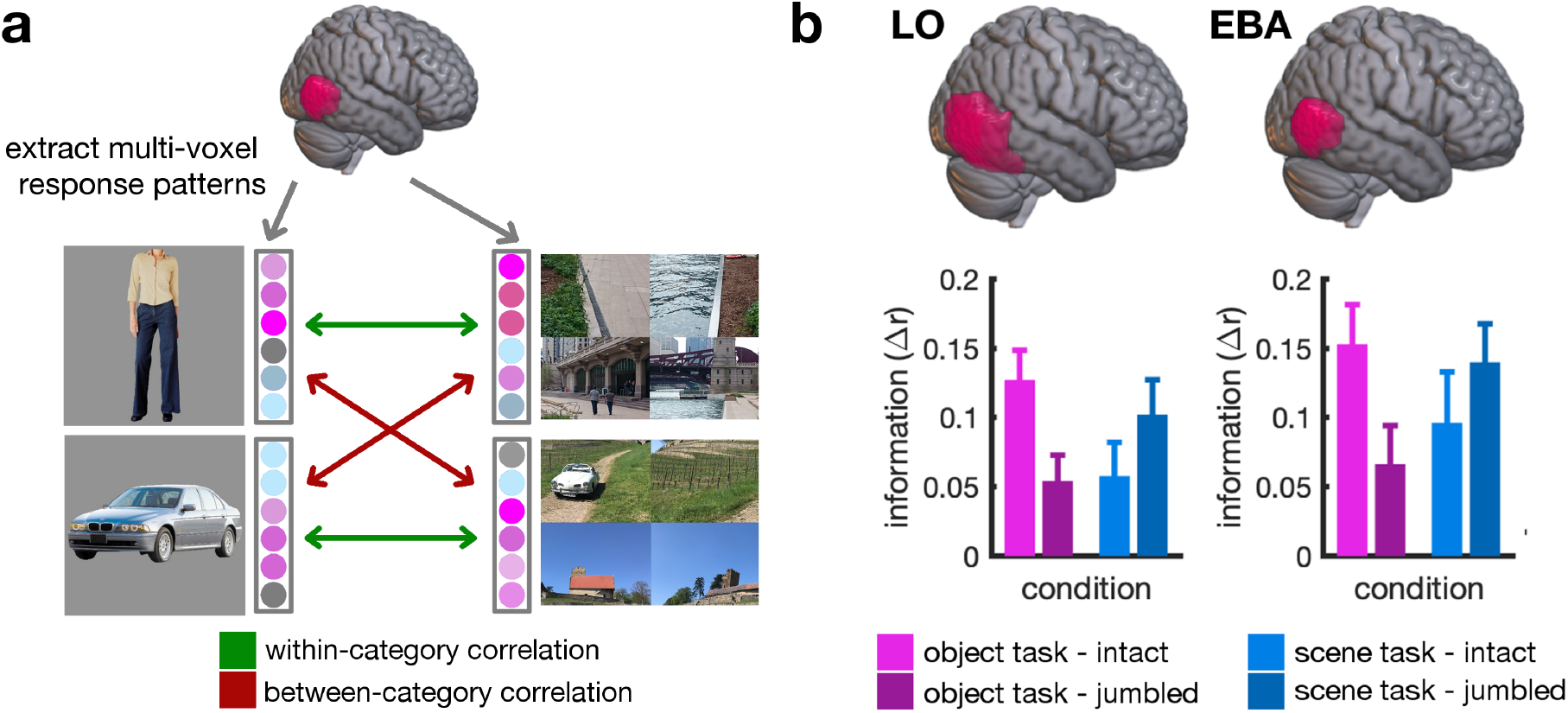
Correlation MVPA logic and results. **a)** To measure object discriminability, we extracted multi-voxel response patterns for each ROI, separately for objects in isolation (from the benchmark localizer) and objects appearing within the scenes (from the main experiment). We then computed within- and between-category correlations. By subtracting the between-category from the within-category correlations, we obtained an index of category information (Δr). **b)** In both LO and EBA, category information was significantly higher for objects that were embedded in intact scenes than for objects embedded in jumbled scenes. However, this was only true when participants performed the object task; when they performed the scene task, no significant difference in object category information was observed when comparing intact and jumbled scenes. For illustration purposes, ROI masks are shown on the right hemisphere of a standard-space template using MRIcroGL (Li et al., 2016); the displayed results are averaged across ROIs in both hemispheres. Error bars represent standard errors of the mean.

To quantify object information, we computed a correlation measure by subtracting correlations between different categories (e.g., person in isolation and car within a scene) from correlations between the same categories (e.g., person in isolation and person within a scene) (Figure 3a). This measure was computed separately for each of the object and scene tasks, the intact and jumbled scenes, and all ROIs.

To test whether multi-voxel response patterns contained any information at all about the object contained in the scenes, we first averaged the correlation measure across all conditions. We then tested whether the average category information was significantly different from zero, separately for each ROI. As expected, people and cars could be reliably discriminated from response patterns in the object-selective LO, t(24)=7.56, p<.001, d=1.51, and body-selective EBA, t(24)=8.00, p<.001, d=1.60, but not from response patterns in EVC, t(24)=0.80, p=.43, d=0.18, or scene-selective OPA, t(24)=0.49, p=.63, d=0.10, and PPA, t(24)=0.70, p=.49, d=0.14.

Given that we only found robust object information in LO and EBA, we only performed further analyses for these two regions (Figure 3b). Data were again analyzed in a 2×2 ANOVA with factors scene structure (intact vs jumbled) and task (object task vs scene task), separately for LO and EBA.

When analyzing the amount of object information contained in LO response patterns, we found a significant interaction between task and scene structure, F(1,24)=5.63, p=.026, η_p_^2^=0.19: When participants performed the object task, object information in LO was more pronounced for objects embedded in intact compared to jumbled scenes, t(24)=2.65, p=.014, d=0.53. This effect was absent when participants performed the scene task, t(24)=1.22, p=.24, d=0.24. A similar interaction effect was found in the EBA, F(1,24)=5.19, p=.032, η_p_^2^=0.18: Object information was again enhanced for intact scenes during the object task, t(24)=2.30, p=.030, d=0.46, but not during the scene task, t(24)=0.92, p=.37, d=0.18. These results demonstrate that coherent scene structure indeed enhances object representations in visual cortex. However, this enhancement depends on the behavioral relevance of the object: When scene category, rather than object category, was task-relevant, no such enhancement was observed.

## 4 Discussion

### 4.1 Coherent scene structure facilitates task-relevant object processing

In this study, we shed light on neural object processing in situations where the object is either embedded within a coherent, intact scene or an incoherent, jumbled scene. Consistent with classical studies (Biederman, 1972; Biederman et al., 1973, 1974), our participants were more accurate and faster in perceiving intact, compared to jumbled scenes, both when performing an object categorization task and a scene categorization task. Our univariate findings are consistent with previous fMRI work (Kaiser et al., 2020a): We replicate the finding that intact scenes yield stronger neural responses than jumbled scenes, across high-level visual cortex and prominently in scene-selective regions. This suggests a widespread sensitivity to typical scene structure in the visual system. Importantly, our current results show that scene structure also matters when it comes to the neural representation of objects within the scene: When analyzing the amount of object information contained in multi-voxel response patterns in object and body-selective visual cortex, we found an enhancement of object information when the objects were embedded within intact scenes, compared to jumbled scenes. Critically, this enhancement only emerged in the object categorization task, suggesting that coherent scene structure facilitates the extraction of object information only when the objects are relevant for current behavioral goals.

### 4.2 Interactions between object and scene processing are mediated by scene structure

Our findings support the view that the scene and object processing pathways are not functionally separate, but that scene information can aid the extraction of object information (Brandmann & Peelen, 2017). Theories of contextual facilitation propose that scene structure is analyzed rapidly, potentially based on coarse low-spatial frequency information (Bar, 2004; Bar et al., 2006). This idea is consistent with the observation that an initial representation of scene meaning – the scene’s “gist” – can be extracted from just a single glance (Greene & Oliva, 2009; Oliva & Torralba, 2006, 2007). Contextual facilitation theories argue that detailed object analysis is facilitated by this more readily available information about scene gist (Bar, 2004; Hochstein & Ahissar, 2002). Informing object analysis through the analysis of coarse scene properties may be particularly useful when perception is challenged by the presence of many distracter items and limited visual exposure. Probing perception with such a challenging task, our study shows that the cross-facilitation between object and scene processing is mediated by the scene’s structural coherence: When the analysis of scene gist is disrupted by jumbling the scene, contextual information cannot amplify object processing in the same way as it can for intact scenes.

The enhanced extraction of object information from the intact scenes suggests that useful information about scene gist is extracted less efficiently from the jumbled scenes. Indeed, the rapid analysis of scene gist depends on our priors about typical scene composition (Csathó et al., 2015; Greene et al., 2015). Neuroimaging studies suggest that the cortical scene processing network is tuned to these priors (Kaiser et al., 2020a; Torralbo et al., 2013), and that the early extraction of properties like the scene’s basic-level category depends on the structural coherence of the scene (Kaiser et al., 2020b). Jumbling is a strong manipulation in the sense that is disrupts multiple aspects of the scene’s spatial coherence at the same time: it disrupts the spatial positioning of individual pieces of information in visual space (Kaiser et al., 2018; Mannion, 2015), the positioning of objects relative to each other (Kaiser et al., 2019; Kaiser & Peelen, 2018), as well as the typical geometry of the scene (Dillon et al., 2018; Spelke & Lee, 2012). Future research is needed to disentangle these different factors, and how much they each contribute to the facilitation of object representation.

Alternatively, one could argue that the jumbling manipulation generates a more general “artificiality” in the stimuli (through the salient borders between quadrants of the jumbled images) that puts additional strain on the visual system. Based on this assertion, one would predict lower responses for jumbled scenes. In previous studies (Kaiser et al., 2020a, 2020b), we have shown that strong effects of scene jumbling are also obtained when introducing similar artificial discontinuities to the typical scenes, suggesting that the degree of image artificiality introduced by the jumbling manipulation alone cannot explain the results.

However, although jumbling is a strong manipulation that conflates multiple factors of scene structure, it preserves critical characteristics of the objects: First, the objects remain completely unaltered across the intact and jumbled scenes. Second, the objects’ absolute positions in visual space were matched across the intact and jumbled scenes. Finally, each object’s local visual context remains constant across the intact and jumbled scenes. These properties allow us to attribute differences in object representations to facilitates effects from cortical scene analysis: If the visual brain would not take global scene context into account and would only analyze the objects in their local visual surroundings, our paradigm should yield comparable results for structurally coherent, intact scenes and incoherent, jumbled scenes.

### 4.2 Attention mediates contextual facilitation effects

Unlike task-relevant objects, task-irrelevant objects were not processed differently as a function of scene coherence. This finding shows that contextual facilitation of object processing is not an automatic process. On the contrary, interactions between the object and scene processing systems seem to be mediated by attention. This observation fits well with previous results from studies on object detection in natural scenes. Compared to task-relevant objects, multi-voxel response patterns in visual cortex contain far less information about unattended objects (Peelen et al., 2009; Peelen & Kastner, 2011). Further, MEG decoding results suggest strong differences in the representation of attended and unattended object categories (Kaiser et al., 2016): Particularly at early stages of processing, within the first 200ms after stimulus onset, the category of unattended objects is represented less accurately. Beyond the visual brain, differences in task demands also affect more widespread activations across the cortex (Cukur et al., 2013; Harel et al., 2014; Hebart et al., 2018; Nastase et al., 2017), potentially causing substantial task-related changes in processing dynamics. One such change may be an alteration of the crosstalk between representations in different visual domains. Our data indeed suggests that the exchange of information between the object and scene processing pathways is not mandatory, but rather constitutes an adaptive mechanism for improving task performance. Under this view, interactions between the scene and object processing pathways may be specifically “switched on” when objects are part of current attentional templates (Battistoni et al., 2017; Peelen & Kastner, 2011). The specific mechanism underlying this adaptive control of the crosstalk between scene and object processing needs further investigation.

How does the apparent importance of attention tie in with previous studies that reported a cross-facilitation between the object and scene-processing systems (Brandmann & Peelen, 2017, 2019)? While these studies did not use object categorization tasks, they still explicitly asked participants to attend to the objects appearing within the scene (either by asking them to memorize them or through one-back tasks). In our scene categorization task, the situation was entirely different, as the objects were completely irrelevant for solving the task. In fact, this orthogonality of object and scene category in our design may have introduced an active suppression of object information when participants performed the scene categorization task. Previous studies suggest that task-irrelevant distracter objects can be suppressed effectively and quickly (Seidl et al., 2012; Hickey et al., 2019). During the scene task, we indeed found numerically better object representations for jumbled scenes. This tentative reversal of the facilitation effect could potentially hint at a more efficient suppression of object information when the object is embedded in a structurally coherent scene. However, as this reversal is not statistically significant in our data and is somewhat susceptible to changes in analysis choices (see Supplementary Information), this assertion is largely speculative at this point. As another interesting observation, object information for the jumbled scenes was comparable between the object and scene tasks, suggesting that attention cannot as efficiently amplify object information when the scene is jumbled. However, some caution needs to be applied when directly comparing representations across the two tasks (rather than comparing differences between conditions), because different task-specific demand characteristics and attentional requirements complicate the interpretation of such comparisons.

### 4.4 Conclusion

In conclusion, our results show that the object and scene processing pathways can interact to facilitate the processing of task-relevant object information embedded in coherent scenes. However, such interactions are not mandatory. They rather seem to be guided by current behavioral goals. Our findings therefore suggest that the visual brain adaptively exploits coherent scene context to resolve object perception in challenging real-world situations.

## Acknowledgements

We thank Sina Schwarze for helping with the stimulus collection. D.K. and R.M.C. are supported by Deutsche Forschungsgemeinschaft (DFG) grants (KA4683/2-1, CI241/1-1, CI241/3-1). G.H. is supported by a PhD fellowship of the Einstein Center for Neurosciences Berlin. R.M.C. is supported by a European Research Council Starting Grant (ERC-2018-StG 803370).

