## Supplementary Information for "Coherent natural scene structure facilitates the extraction of task-relevant object information in visual cortex"

Supplementary Contents | Page

*Figure S1. Univariate results – category-selective responses* 2

*Figure S2. Univariate results separately for both hemispheres* 3

*Figure S3. MVPA results separately for both hemispheres* 4

*Figure S4. MVPA results with alternative analysis routines* 5

*Table S1. Descriptive statistics – behavior* 6

*Table S2. Descriptive statistics – univariate analysis* 7

*Table S3. Descriptive statistics – MVPA* 8

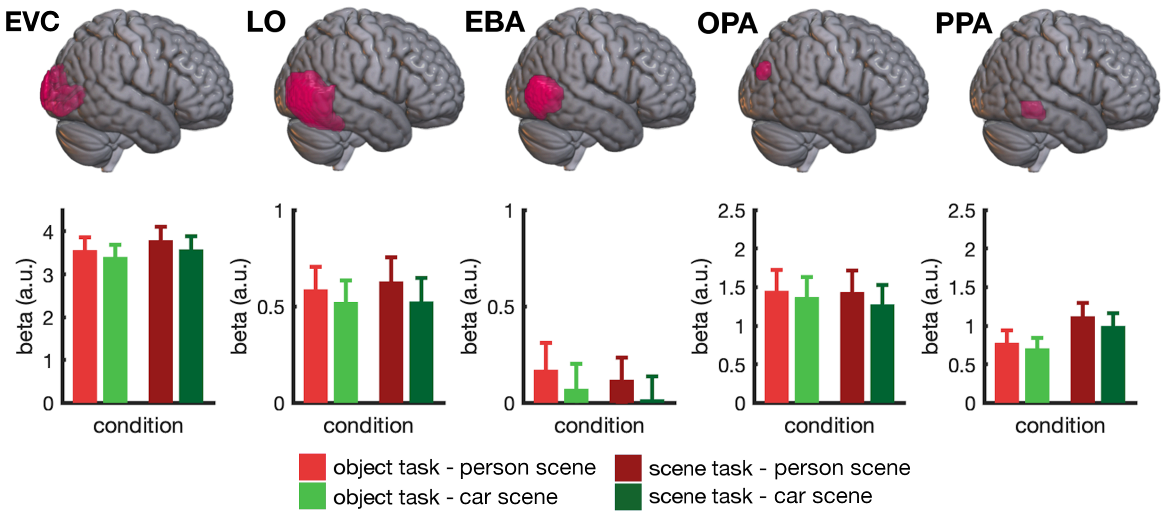


***Figure S1. Univariate results – category-selective responses.*** *Across all regions, we found stronger responses to scenes that contained a person than to scenes that contained a car (collapsed across intact and jumbled scenes), main effect of category across ROIs, F(1,24)=8.50, p=0.008,* $\eta_p^2$*=0.62. This general bias towards person-scenes was not modulated by participants' current task, category × task interaction, F(1,24)=0.30, p=0.59,* $\eta_p^2$*=0.01. For illustration purposes, ROI masks are shown on the right hemisphere of a standard-space template using MRIcroGL (Li et al., 2016); the displayed results are averaged across ROIs in both hemispheres. Error bars represent standard errors of the mean.*

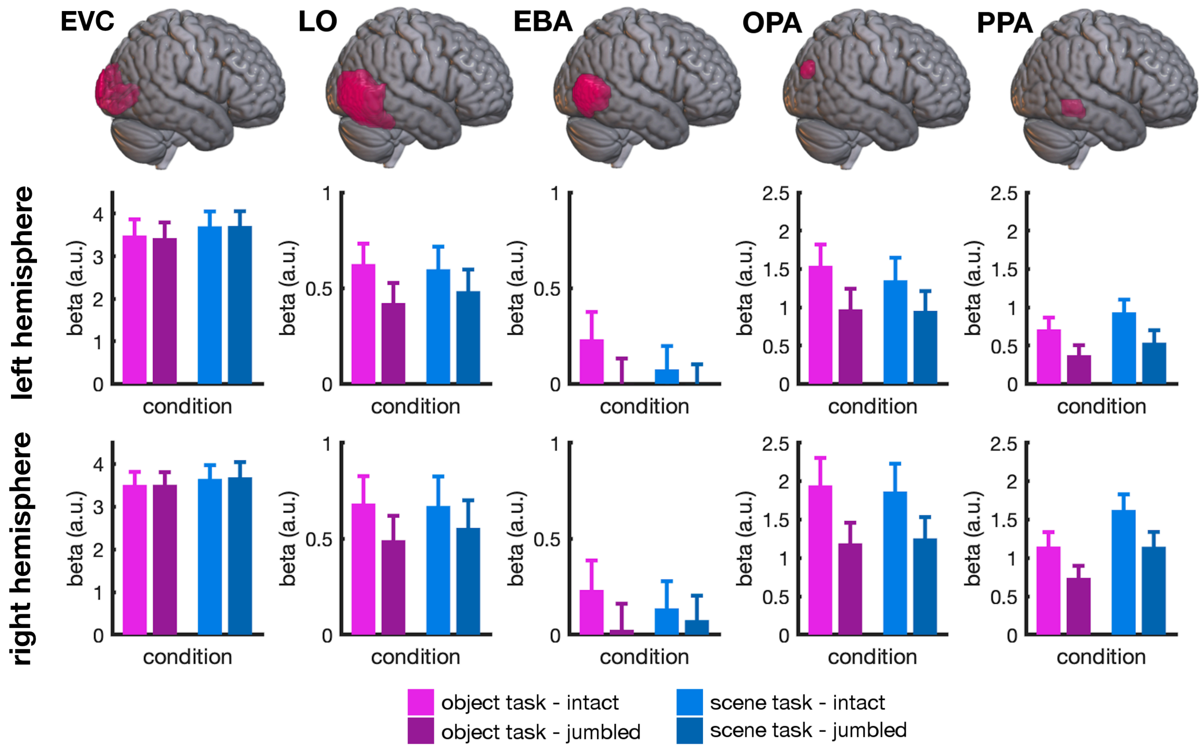


***Figure S2. Univariate results separate for both hemispheres.*** *Results for the left and right hemispheres were highly similar and closely resembled the results across both hemispheres (Figure 2). No qualitative difference between hemispheres was found, as indicated by non-significant main effects and interactions in all ROIs. The only exception was a hemisphere × scene structure interaction in OPA, $F(1,24)=6.31$, $p=0.019$, $\eta_p^2=0.21$, with a stronger effect of scene structure in the right hemisphere. For illustration purposes, ROI masks are shown on the right hemisphere of a standard-space template using MRIcroGL (Li et al., 2016). Error bars represent standard errors of the mean.*

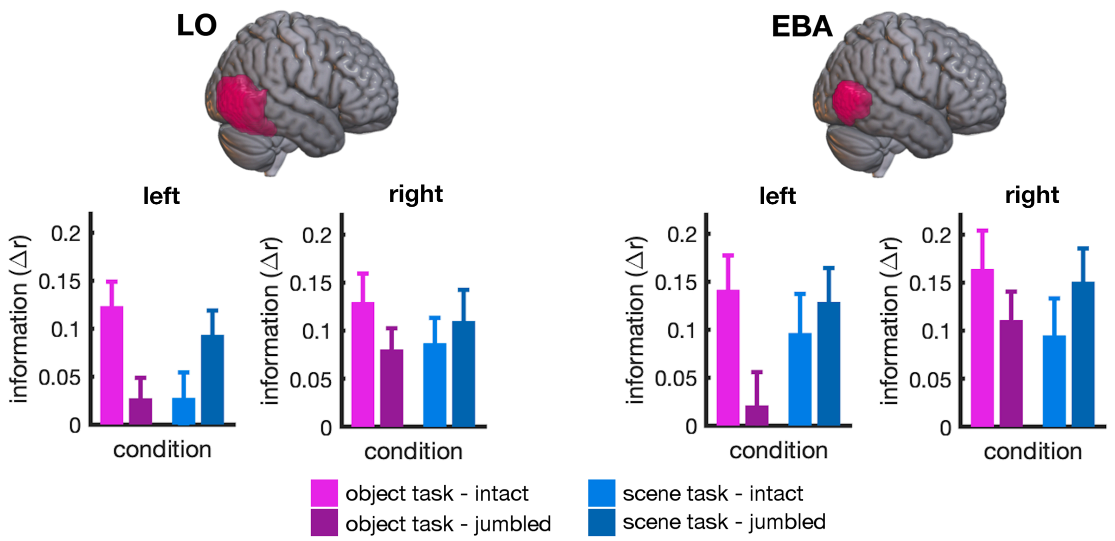


***Figure S3. MVPA results separately for both hemispheres.*** *Results for the left and right hemispheres closely resembled the results across both hemispheres (Figure 3). No qualitative difference between hemispheres was found, all interactions with hemisphere, LO: $F(1,24)<2.03$, $p>0.16$, $\eta_p^2<0.08$, EBA: $F(1,24)<2.49$, $p>0.12$, $\eta_p^2<0.10$. In LO, category information was generally stronger in the right-hemispheric than in the left-hemispheric ROI, main effect of hemisphere, $F(1,24)=5.34$, $p=0.030$, $\eta_p^2=0.18$. For illustration purposes, ROI masks are shown on the right hemisphere of a standard-space template using MRIcroGL (Li et al., 2016). Error bars represent standard errors of the mean.*

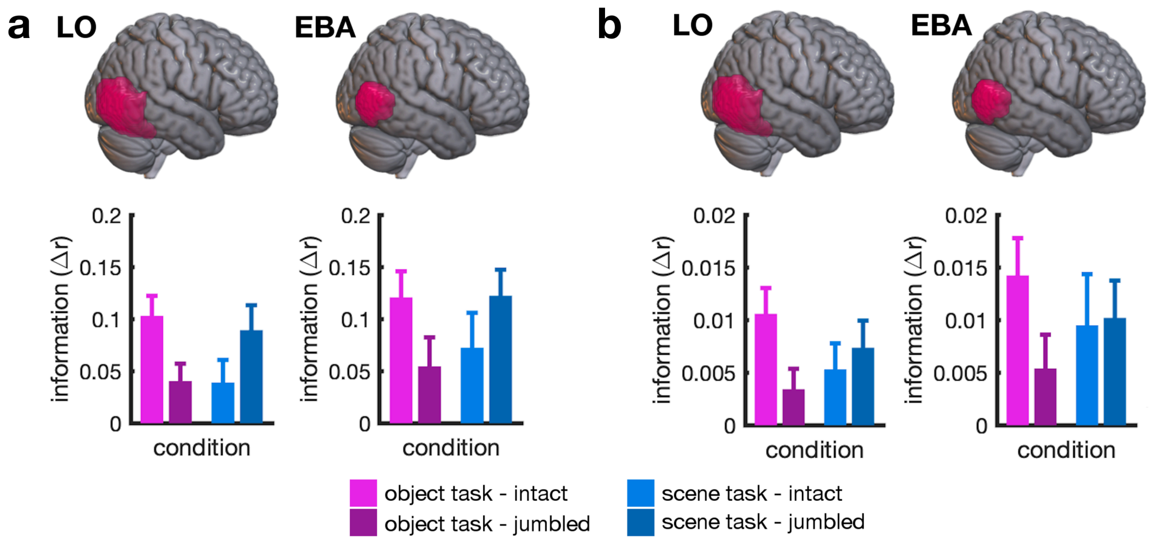


***Figure S4. MVPA results with alternative analysis routines. a)*** *Category information in LO and EBA for intact and jumbled scenes across the two tasks, computed using Spearman correlations instead of Pearson correlations. As in the main analysis (Figure 3), an interaction between task and scene structure was observed for both regions, LO: $F(1,24)=6.29$, $p=0.019$, $\eta_p^2=0.21$, EBA: $F(1,24)=5.10$, $p=0.033$, $\eta_p^2=0.18$.* ***b)*** *Category information, as in (a), but computed without removing the voxel-wise mean activation across conditions. Again, an interaction effect was observed for LO: $F(1,24)=4.48$, $p=0.045$, $\eta_p^2=0.16$, but did not reach significance in EBA: $F(1,24)=2.33$, $p=0.14$, $\eta_p^2=0.09$. For illustration purposes, ROI masks are shown on the right hemisphere of a standard-space template using MRIcroGL (Li et al., 2016); the displayed results are averaged across ROIs in both hemispheres. Error bars represent standard errors of the mean.*

***Table S1. Descriptive statistics – behavior.*** *Means (M) and standard errors (SE) for the accuracies (in % correct) and response times (in ms), as shown in Figure 1.*

| | ***Object Task Intact*** | ***Object Task Scrambled*** | ***Scene Task Intact*** | ***Scene Task Scrambled*** |
|---|---|---|---|---|
| ***Accuracy*** | M=77.1<br>SE=1.6 | M=70.5<br>SE=1.5 | M=76.7<br>SE=1.4 | M=73.7<br>SE=1.5 |
| ***Response Time*** | M=717<br>SE=16 | M=731<br>SE=17 | M=715<br>SE=15 | M=736<br>SE=17 |

***Table S2. Descriptive statistics – univariate analysis.*** *Means (M) and standard errors (SE) for univariate activations in each ROI, as shown in Figure 2.*

| ***ROI*** | ***Object Task Intact*** | ***Object Task Scrambled*** | ***Scene Task Intact*** | ***Scene Task Scrambled*** |
|---|---|---|---|---|
| ***EVC*** | M=3.50<br>SE=0.30 | M=3.47<br>SE=0.29 | M=3.68<br>SE=0.30 | M=3.70<br>SE=0.32 |
| ***LO*** | M=0.65<br>SE=0.12 | M=0.46<br>SE=0.11 | M=0.63<br>SE=0.13 | M=0.52<br>SE=0.12 |
| ***EBA*** | M=0.23<br>SE=0.14 | M=0.01<br>SE=0.13 | M=0.11<br>SE=0.12 | M=0.03<br>SE=0.11 |
| ***OPA*** | M=1.74<br>SE=0.29 | M=1.08<br>SE=0.24 | M=1.61<br>SE=0.30 | M=1.10<br>SE=0.24 |
| ***PPA*** | M=0.93<br>SE=0.16 | M=0.56<br>SE=0.13 | M=1.28<br>SE=0.17 | M=0.84<br>SE=0.17 |

***Table S3. Descriptive statistics – MVPA.*** *Means (M) and standard errors (SE) for object discriminability (difference of within- and between-category correlations) in each ROI, as shown in Figure 3.*

| ***ROI*** | ***Object Task Intact*** | ***Object Task Scrambled*** | ***Scene Task Intact*** | ***Scene Task Scrambled*** |
|---|---|---|---|---|
| ***LO*** | M=0.13<br>SE=0.02 | M=0.05<br>SE=0.02 | M=0.06<br>SE=0.02 | M=0.10<br>SE=0.03 |
| ***EBA*** | M=0.15<br>SE=0.03 | M=0.07<br>SE=0.03 | M=0.10<br>SE=0.04 | M=0.14<br>SE=0.03 |
